# Early transcriptional signatures of MeCP2 positive and negative cells in Rett syndrome

**DOI:** 10.1101/2025.06.26.661761

**Authors:** Yan Li, Ashley G. Anderson, Guangtong Qi, Sih-Rong Wu, Jean-Pierre Revelli, Zhandong Liu, Huda Y. Zoghbi

## Abstract

Rett syndrome (RTT) is an X-linked neurological disorder caused by *MECP2* mutations. Like other X-linked disorders, RTT patients have sex-specific differences in clinical presentation due to distinct cellular environments, where females have ∼50% of cells expressing either a mutant or wild-type copy of *MECP2* (mosaic) and males have 100% of cells expressing a mutant *MECP2* (non-mosaic). Typical RTT females have a short window of normal early development until ∼6-18 months, followed by regression and progressive decline, whereas neonatal encephalopathy is more likely in RTT males. How these sex-specific differences in cellular context contribute molecularly to RTT pathogenesis, particularly in the presymptomatic stages of RTT females, remains poorly understood. Here, we profiled the hippocampal transcriptomes of female (*Mecp2*^+/-^) and male (*Mecp2*^-/y^) RTT mice at early timepoints using both bulk and single-nucleus RNA-seq, including sorted MeCP2 positive (MeCP2+) and MeCP2 negative (MeCP2-) neurons in female mice. We identified a core disease signature consisting of 12 genes consistently dysregulated only in MeCP2-cells across RTT models. Moreover, we uncovered non-cell-autonomous effects exclusively in female MeCP2+ excitatory neurons, but not inhibitory neurons, suggesting excitatory circuits are more vulnerable early in the mosaic RTT environment. The single-nuclei data also revealed that a previously underappreciated MeCP2-interneuron subtype had the most transcriptional dysregulation in both male and female RTT hippocampi. Together, these data highlight the different effects of MeCP2 loss on excitatory and inhibitory circuits between the mosaic and non-mosaic environment that appear early in RTT pathogenesis.

## Introduction

Rett syndrome (RTT, OMIM:312750) is a severe neurological disorder caused by loss-of-function mutations in the X-linked gene methyl-CpG binding protein 2 (*MECP2)* (*1*). Classic RTT predominantly affects females and is characterized by a period of normal development until ∼6-18 months, followed by developmental regression. This regression is marked by loss of acquired motor, language, and social skills, development of stereotyped hand movements, irregular breathing, and intellectual disability (*2*). A key feature of RTT is the unique mosaic cellular environment in females, where random X-chromosome inactivation (XCI) results in approximately half of cells expressing either a mutant or wild-type copy of *MECP2* (*3*). This mosaicism fundamentally shapes disease presentation, as shown by the correlation between XCI skewing patterns and phenotypic severity (*4*, *5*). In contrast, males with germline *MECP2* mutations, where all cells express mutant *MECP2*, typically present with neonatal encephalopathy unless the mutation is relatively mild (*6–9*). The difference between female (mosaic) and male (non-mosaic) RTT raises important questions about how cellular mosaicism influences neuronal dysfunction in RTT pathogenesis and whether distinct molecular pathways emerge in these different cellular contexts.

*MECP2* encodes a methyl cytosine binding protein (MeCP2) that is highly abundant in postnatal neurons and functions as a transcriptional regulator, whose dysfunction causes thousands of genes to be up-or down-regulated (*10–15*). While studies have shown that animal models of *Mecp2* loss-of-function mutations effectively recapitulate the human disorder (*16–18*), there are important differences between mosaic female *Mecp2^+/-^* and non-mosaic male *Mecp2^-/y^* mice. Female *Mecp2^+/-^* mice show a progressive disease course that parallels classic RTT, while male *Mecp2^-/y^* mice develop severe early phenotypes and die at a young age (8-12 weeks) (*19*, *20*). Male mouse models provide a valuable system for studying the direct molecular consequences of complete MeCP2 loss and have been the predominant model for most molecular studies in the field. Far fewer studies have examined the molecular changes in female RTT models (*21–24*), leaving a gap in our understanding of RTT disease progression in the mosaic cellular environment.

While most molecular studies are performed at symptomatic stages of the disease, recent studies from our lab have highlighted the importance of studying presymptomatic timepoints in RTT (*25*, *26*). First, training of female RTT mice in the presymptomatic but not in the symptomatic phase can significantly delay symptom onset, suggesting a critical window for such an intervention to overcome MeCP2 loss (*26*). Second, transcriptional dysregulation precedes functional changes by several weeks in the hippocampus of adult male mice after acute deletion of *Mecp2*, showing that early transcriptional dynamics drive disease pathogenesis (*25*). These observations led us to ask a series of questions: do RTT females show pre-symptomatic transcriptional changes? If so, are these changes shared between male and female RTT models? And finally, are these molecular changes specific to distinct cell-types, including MeCP2+ and MeCP2-cells in the mosaic context?

To answer these questions, we designed experiments using both bulk RNA sequencing (RNA-seq) and single-nucleus RNA sequencing (snRNA-seq) to molecularly profile the hippocampus of male (NULL, *Mecp2^-/y^*) and female (HET, *Mecp2^+/-^*) RTT mice. We examined two early timepoints in disease progression (P28 – 4 weeks and P45 – 6.5 weeks). At 4 weeks, *Mecp2* transcript and protein levels become stable in the postnatal brain, and at 6.5 weeks female RTT mice have no measurable behavioral deficits. We focused our transcriptional analysis on the hippocampus, a region where disruption of excitatory and inhibitory (E/I) balance, accompanied by learning and memory deficits, is among the earliest and most pronounced phenotypes in both male and female RTT mice (*19*, *27–32*).

Using bulk RNA-seq, we identified an early molecular disease signature shared across both male and female RTT mouse models, independent of timepoint. Using snRNA-seq, we profiled >120,000 nuclei, including sorted MeCP2+ and MeCP2-neurons from female RTT hippocampus, and revealed this early disease signature is specific to MeCP2-neurons in the mosaic brain. Our single-nucleus analyses gave us the resolution to uncover shared and unique cell-type specific molecular changes in the mosaic and non-mosaic RTT hippocampus. Notably, we found widespread transcriptional changes in MeCP2-and, surprisingly, MeCP2+ female RTT hippocampal excitatory neurons, suggesting excitatory neurons in the mosaic condition are highly sensitive to the altered circuit environment early in disease pathogenesis. We also identified an interneuron subpopulation (with high *Chrm2* expression) that showed a prominent transcriptional response to direct loss of MeCP2 in both male and female RTT. Together, this study uncovered an early molecular disease signature in both male and female RTT that is MeCP2 dependent and not secondary to broader circuit dysfunction. It also provides insight into the distinct differences of cell-autonomous and non-cell-autonomous changes in excitatory and inhibitory neurons of RTT females. These data provide important neurobiological insights into the cellular and molecular drivers of pathogenesis in RTT, and that are potentially relevant to other X-linked disorders.

## Results

### Early disease signature in the RTT hippocampus

Since male and female RTT mice have distinct time courses for disease progression, we examined whether there is a shared disease signature that is consistent between sexes and can be detected early in disease pathogenesis. We first compared the transcriptional changes in the female and male RTT hippocampus using bulk RNA-seq in a mouse model with a null *Mecp2* allele (*Mecp2^tm1.1Bird/J^*) (*16*), which produces no MeCP2 protein and shows hippocampal phenotypes at 6 weeks in NULL mice and at 12 weeks in HET mice (*16*, *19*). To avoid the confounds of phenotypic and secondary changes between male and female RTT mice, we used early timepoints in disease course (P28 and P45) and examined differentially expressed genes (DEGs) between genotypes within each sex and timepoint (**Figure 1A**). Principal component analysis (PCA) of all samples together shows that samples clustered primarily by sex (**Figure S1A**). Male samples showed clear separation by timepoint and genotype, while female samples were more interspersed (**Figure S1A**). We found that NULL mice had ∼3X more DEGs at each timepoint compared to HET mice (**Figure 1B**). The magnitude of expression level changes was also greater in NULL compared to HET mice at both timepoints (**Figure 1B**). In NULL samples, the number of both up- and down-regulated DEGs increased progressively from P28 (313 down, 298 up; total 611 DEGs) to P45 (370 down, 437 up; total 807 DEGs). Despite showing no measurable phenotypes at these early timepoints, female HET mice exhibited detectable transcriptional changes at P28 (21 down, 46 up; total 67 DEGs) and P45 (49 down, 46 up; total 95 DEGs).

**Figure 1.**
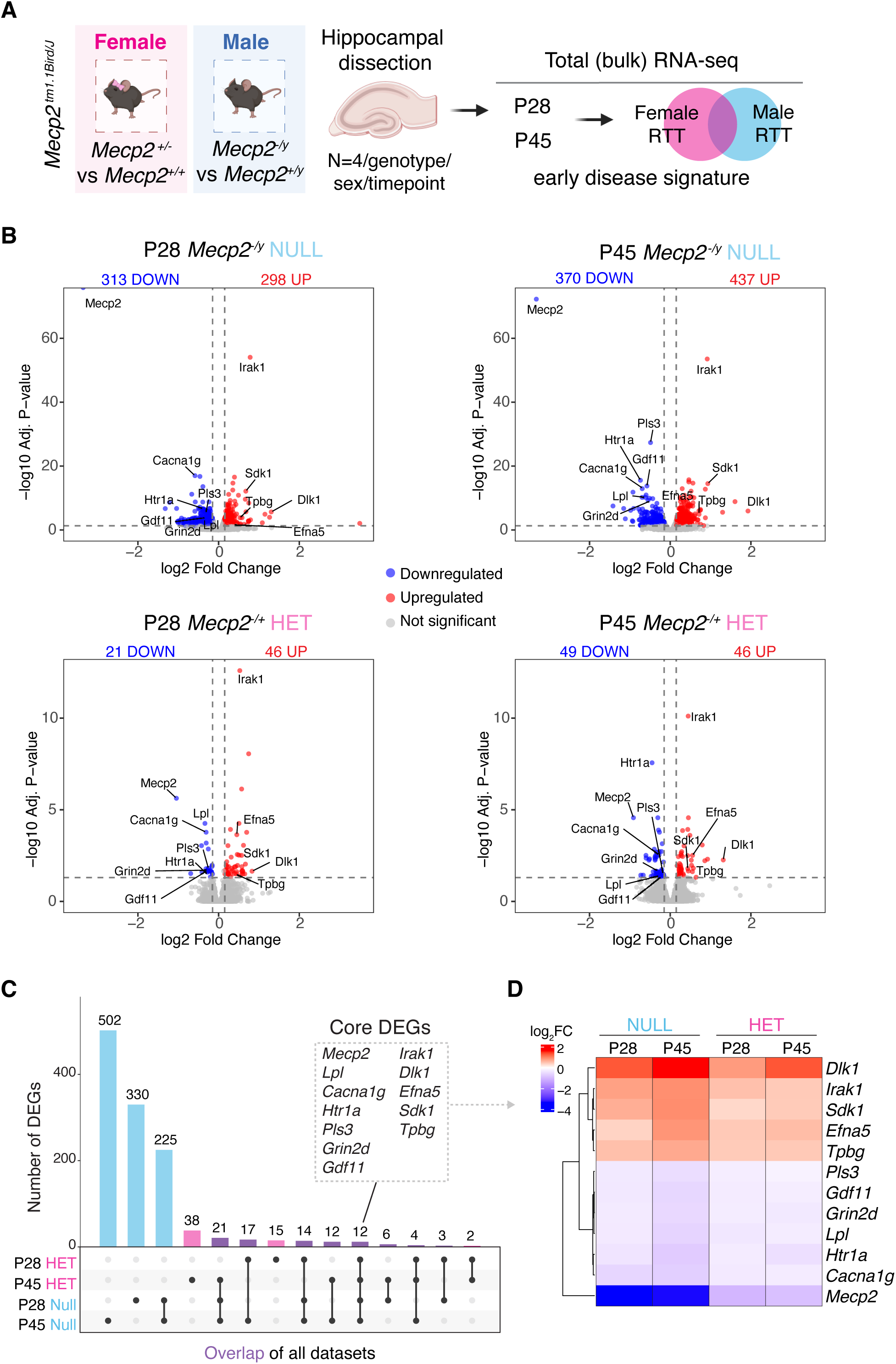
Bulk RNA-seq of hippocampal tissue in male and female RTT mice at pre- and early-symptomatic timepoints. **A**) Schematic of experimental design using hippocampal tissue from male and female RTT mice along with sex-specific controls at two distinct timepoints (P28 and P45) to examine early transcriptional disease signatures via bulk RNA-seq. Created with BioRender.com. **B**) Volcano plots showing significant differentially expressed genes (DEGs) that are upregulated (red) or downregulated (blue) at both timepoints in male NULL (top panels) or female HET (bottom panels) compared to their sex- and age-matched control WT samples using a |log_2_FC|>0.15 and adjusted p-value <0.05 cutoff for all comparisons. **C**) Upset plot showing the overlap of DEGs across timepoints and genotypes with 12 core DEGs highlighted. **D**) Heatmap showing the log_2_FC of core RTT DEGs at each timepoint within NULL and HET samples.

Overlapping the DEGs across both genotypes and timepoints, we found 12 DEGs (termed “core RTT DEGs”), including *Mecp2*, that were persistently dysregulated in HET and NULL samples in the same direction, indicative of an early disease molecular signature across RTT models (**Figure 1C**). Five genes were consistently upregulated (*Dlk1*, *Irak1*, *Sdk1*, *Efna5*, and Tpbg) and 7 genes were consistently downregulated (*Pls3*, *Gdf11*, *Grin2d*, *Lpl*, *Htr1a*, *Cacna1g*, *Mecp2*) (**Figure 1D**).

Interestingly, 7 out of 12 genes (*Cacna1g*, *Htr1a*, *Efna5*, *Tpbg*, *Grin2d*, *Sdk1*, and *Pls3*) are involved in synapse-related gene ontology categories (data not shown). Moreover, the magnitude of change in core RTT DEGs increased as the animals aged, particularly within NULL mice (**Figure 1D**).

Our lab has previously shown that *Gdf11* is sensitive to *Mecp2* level changes (*33*), which we confirm can be detected as early as 4 weeks in both male NULL and female HET tissue. The biological significance of these core RTT DEGs is supported by their significant overlap with our previous study examining the molecular cascade following the acute loss of MeCP2 within the hippocampus of adult male mice using the *Mecp2^tm1.1Jae^* floxed allele (adult KO) (*17*, *25*). We found that 11 of our core RTT DEGs (all but *Irak1*) overlapped in the same direction with DEGs from acute loss of MeCP2 between 4-8 weeks when MeCP2 levels were stably depleted (17.1-fold enrichment, p < 1.0 × 10⁻¹⁰, hypergeometric test) (**Figure S1B**). The unique observation that *Irak1* is only changed in the constitutive knockout but not after acute *Mecp2* loss in adult mice suggests that changes in *Irak1* may be due to either a developmental effect of *Mecp2* on this gene or a local effect of the engineered *Mecp2* allele, given that *Irak1* is located only 3 kilobases distal to the replacement cassette (*16*, *34*).The magnitude of change in expression also progressively increased over time for core RTT DEGs in the adult KO datasets (**Figure S1C**). Together, these data identify a shared core molecular disease signature in both female and male RTT models early in the disease course, before female RTT mice develop neurological symptoms (*19*), suggesting these transcriptional changes represent an important step in disease pathogenesis rather than being secondary to the RTT phenotype.

### Transcriptional changes in male and female RTT hippocampi are restricted to mature cell-types

MeCP2 is broadly expressed in various cell types throughout the brain, but its protein levels vary among different cell populations (*35–37*). Furthermore, cell-type specific deletion of *Mecp2* results in distinct phenotypic patterns in mouse models (*38–41*). To investigate whether different cell types exhibit differential sensitivity to MeCP2 loss, we analyzed gene expression changes using single nucleus RNA-sequencing (snRNA-seq) in the hippocampus of both male and female RTT mice. We isolated hippocampal nuclei via fluorescence-activated nuclei sorting (FANS) from *Mecp2^-/y^*(NULL) and *Mecp2^+/-^* (HET) animals at 6.5 weeks, matching the timepoint in our bulk RNA-sequencing dataset (**Figure 2A**). We targeted 10,000 nuclei per sample (N=3/group, 12 samples total) and 98,266 nuclei passed quality control criteria (**Figure S2A**). Importantly, we did not find differences in total RNA count across genotypes (**Figure S2A**). In the snRNA-seq dataset, we observed more reads mapping to the *Mecp2* transcripts in NULL and HET samples. However, we confirmed that none of the reads mapped to the deleted exons 3-4 region in the *Mecp2^tm1.1Bird^* allele, indicating the absence of functional *Mecp2* transcript (**Figure S2B**). Unsupervised clustering detected a total of 34 clusters (**Figure 2B)**. We annotated the clusters by comparing their top unique marker genes to the known marker genes of hippocampal cell types from previous single-cell studies (**Figure 2B, S2D)** (*42*, *43*). We did not observe differences in cell-type composition across genotypes (**Figure 2C, S2C**), validating previous findings that loss of MeCP2 neither alters cell identity nor leads to cell death (*44*). To identify cell-type specific DEGs, we performed differential expression analysis within each cluster comparing female and male RTT models to their respective sex and littermate matched controls (*Mecp2^+/-^* vs *Mecp2^+/+^* females or *Mecp2^-/y^* vs *Mecp2^+/y^* males) (**Figure 2D**). Both upregulated and downregulated single-nuclei DEGs (snDEGs) were enriched in multiple cell types in male and female RTT mice, including excitatory hippocampal neuronal clusters (DG, CA1, CA2, CA3), distinct interneuron (IN) clusters (INs^CHRM2^, INs^SST/PVAL^), and specific glial clusters (oligodendrocytes and astrocytes) (**Figure 2D**). We detected sex-specific DEG signatures in two cell types: pericytes (cluster 26) were specific to NULL samples, while excitatory neuronal subtype 1 (Ex. N-1, cluster 15) was specific to HET samples. Notably, immature cell-types, such as oligodendrocyte precursor cells (OPCs) or neuroblasts, did not have many transcriptional changes, highlighting the important role of MeCP2 in mature neuronal cell types (**Figure 2D**) (*17*, *25*). Together, these results demonstrate that the most transcriptional changes occur in mature hippocampal cell types in both males and females.

**Figure 2.**
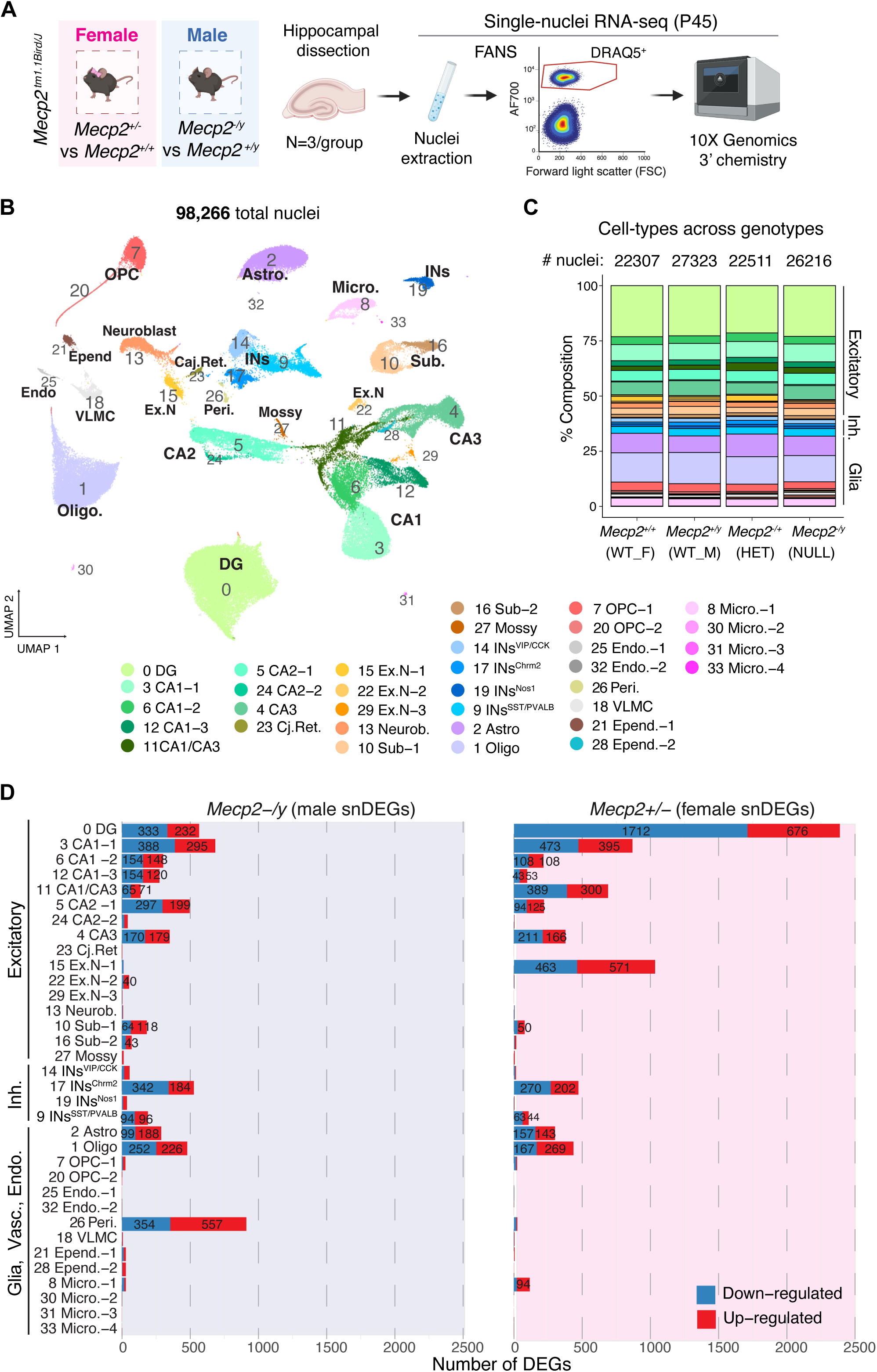
**Single-nucleus RNA-seq of hippocampus from male and female RTT mice**. **A**) Schematic of experimental design of single-nucleus RNA-seq using hippocampus from male and female RTT mice at P45 timepoint. Created with BioRender.com. **B**) UMAP of all hippocampal nuclei passing quality control across genotypes and annotated by cell type. **C**) Bar graph showing the percent cell type composition across each genotype and counts of total nuclei (no significant changes in cell type proportion, see Methods). **D**) Bar graphs showing the number of differentially expressed genes (|log_2_FC|>0.15 and adjusted p-value <0.05) either upregulated or downregulated within each cluster in male (left) and female (right) RTT cell types.

### Comparison of cell-type specific transcriptional changes between male and female RTT models

Since females with mosaic *Mecp2* loss show slower disease progression than males with complete *Mecp2* loss, previous studies comparing male and female models have usually been conducted at different ages. Here, we identify shared disease signatures between age-matched males and females at cell type resolution. We examined the shared molecular signatures of NULL and HET samples within each cell-type by overlapping the snDEGs within each cluster (**Figure 3A**). We found that most snDEGs overlapping within each cluster had concordant gene expression directionality between male and female RTT models. We also found the magnitude of shared DEGs correlated between HET and NULL samples within nearly all clusters, particularly the excitatory neuronal clusters within the CA1, CA2, and CA3 regions (**Figure 3B**). DG excitatory neurons were the least concordant excitatory neuron population (**Figure 3B**). Surprisingly, oligodendrocytes had the highest discordance, with 72% of shared DEGs (97 out of 134) showing opposing regulation patterns in male and female RTT oligodendrocytes (**Figure 3C**). Pathway enrichment analysis of these 97 oligodendrocyte DEGs showed enrichment for GO terms such as synapse organization and synaptic signaling pathways (**Figure 3D**), suggesting oligodendrocyte might be sensitive to the cellular environment and respond differently in the mosaic brain.

**Figure 3.**
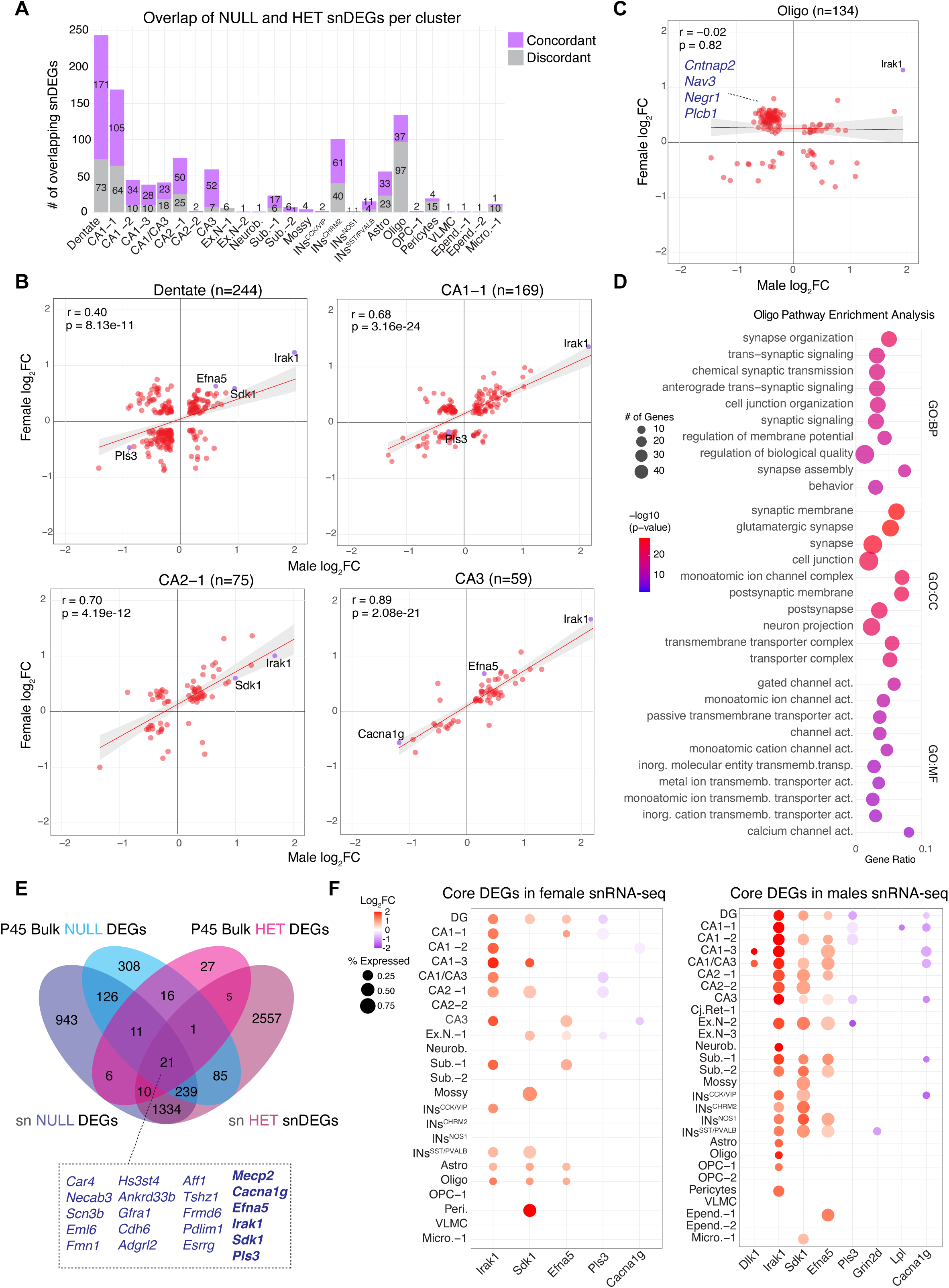
Shared and unique transcriptional signatures across cell-types in hippocampus of male and female RTT mice. **A**) The number of snDEGs overlapping between female and male RTT samples across all clusters. **B**) Correlation of log_2_FC values in shared DEGs between male and female RTT samples in the dentate (DG), CA1-1, CA2-1, and CA3 clusters. **C**) Correlation of log_2_FC values in shared male and female DEGs in the oligodendrocytes and (**D**) the GO enrichment analysis of genes that are upregulated in female but downregulated in male oligodendrocytes. **E**) Venn diagram showing the overlap of bulk RNA-seq DEGs and unique single-nucleus DEGs between male (blue) or female (pink) RTT samples. **F**) Bubble plot showing core DEGs that are also snDEGs across distinct cell types in female (left plot) and male (right plot) RTT hippocampus. Only clusters with snDEGs detected are shown on the plots.

### Core molecular signature is preserved in single-nuclei data across RTT models

We were surprised to find significantly more snDEGs across cell types in female RTT mice compared to males, unlike our bulk RNA-seq results. To compare the bulk and single-nucleus molecular signatures, we overlapped snDEGs and bulk DEGs in both female and male RTT mice (**Figure 3E, Figure S3A, S3C**). We found that ∼50% of bulk DEGs from NULL male mice overlapped with both male snDEGs (**Figure S3B)** and female snDEGs, while ∼40% of bulk female DEGs overlapped with female snDEGs (**Figure S3D**). To better understand whether the changes in snDEGs in the female HET animals represented a true disease signature, we correlated the snDEGs of both female and male RTT animals with the bulk RNA-seq DEGs from male NULL tissue (P45 dataset) and found a significant, positive Pearson correlation between both male (r = 0.619, p = 4.402042e−136) and female (r = 0.565, p = 2.405773e−76) snDEGs with bulk RNA-seq DEGs (**Figure S3E-F**). We also found that most snDEGs were unique to only one cluster (**Figure S3G**) and snDEGs that appeared in more than three clusters (>3) showed a higher percentage overlap with bulk RNA-seq DEGs (**Figure S3G**). These data suggest that snRNA-seq might be more sensitive in capturing early transcriptional changes particularly in female RTT tissue, as cell-type specific changes might be masked in bulk samples.

We next examined the core RTT DEGs in snRNA-seq data, where we captured transcripts of nine of them (all except for *Gdf11*, *Htr1a, and Tpbg).* This is likely due to low capture rate in general of snRNA-seq where ∼30% of transcripts are captured in any given cell (**Figure S2E**) (*45*); however, we did detect 6 core RTT DEGs (including *Mecp2*) in our snRNA-seq data (**Figure 3E, F**). These snDEGs were differentially expressed in the same direction as in the bulk dataset. Importantly, core RTT DEGs identified from our bulk RNA-seq data did not have a strong cell-type specific expression pattern, except for *Grin2d*, an N-methyl D-aspartate receptor (NMDAR) subunit known to be enriched in interneurons (**Figure S2E**). Male RTT snRNA-seq samples had more core DEGs in neurons compared to females, primarily in excitatory neurons of DG, CA1, CA2, and CA3 clusters. Eight core DEGs (*Dlk1*, *Irak1*, *Sdk1*, *Efna5*, *Pls3*, *Grin2d*, *Lpl*, and *Cacna1g*) were differentially expressed in male excitatory and inhibitory neuronal clusters compared to only 5 core DEGs (*Irak1*, *Sdk1*, *Efna5*, *Pls3*, and *Cacna1g*) in females. Interestingly, females had more core DEGs upregulated in glial populations (*Irak1*, *Sdk1*, and *Efna5*) compared to males (*Irak1*) (**Figure 3F**). These cell-type specific changes underscore the importance of studying disease signatures using multiple transcriptomic approaches. To further investigate the effects of female mosaicism, we examined the transcriptional changes of MeCP2+ and MeCP2-neurons separately.

### Single-nuclei analysis of MeCP2+ and MeCP2-neurons in mosaic RTT females

We used an antibody-based sorting strategy to identify MeCP2+ and MeCP2-nuclei in the hippocampus of female HET and WT mice at the same timepoint (P45) as the previous bulk and snRNA-seq datasets (**Figure 4A**). Nuclei were sorted into the following three groups for downstream analyses: 1) WT (*Mecp2^+/+^*, MeCP2+ sorted), 2) HET^POS^ (*Mecp2* ^+/-^, MeCP2+ sorted), and 3) HET^NEG^ (*Mecp2* ^+/-^, MeCP2-sorted). Using similar quality control parameters as the first dataset (**Figure S4A**), we analyzed 22,289 nuclei (∼7-8K nuclei per group) and found that unsupervised clustering separated the nuclei by cell types (**Figure 4B, Figure S4C**). We observed an increase in glial cell types in the HET^NEG^ group compared to WT and HET^POS^ groups (**Figure 4C**). This is due to MeCP2 being more abundant in neurons compared to glia (*36*) and glial populations predominantly fell into the negative gate for HET^NEG^ group when using NULL cells (from male) as the control. As a result, we captured more glial populations in the HET^NEG^ group. Therefore, we subset nuclei from neuronal clusters only and, finding no neuronal cell type composition differences across groups (**Figure 4D, Figure S4B**), focused our downstream analyses specifically on neuronal cell types.

**Figure 4.**
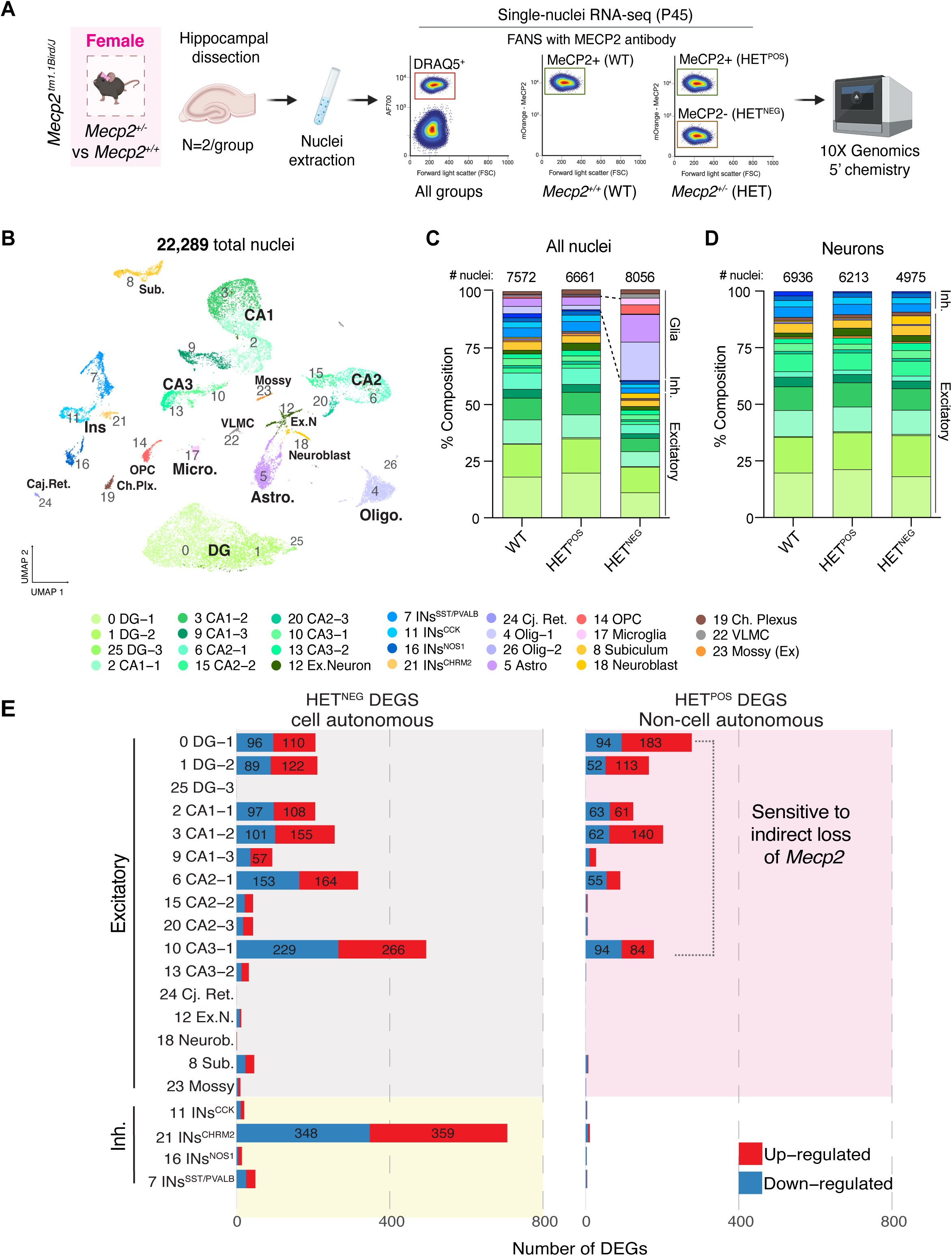
Single-nucleus RNA-seq of MeCP2+ and MeCP2-cells within the mosaic hippocampus of RTT female mice. **A**) Schematic of the experimental design using FANS to isolate nuclei from female wildtype (WT) and *Mecp2^+/-^ (*HET) mice and then using an anti-MeCP2-PE conjugated antibody to capture MeCP2+ (WT, HET^POS^) and MeCP2-(HET^NEG^ only) cells. We used the 10X 5’ chemistry kit to capture nuclei and sequence transcripts. Created with BioRender.com. **B**) UMAP of all female WT, HET^POS^, and HET^NEG^ nuclei that passed quality control parameters like Figure 2. **C**) The percent composition within all three samples across all cell types (non-neuronal clusters 4, 14, 17, 22 showed different proportions in different groups) and (**D**) neurons specifically. **E**) Bar charts showing the number of differentially expressed genes (|log_2_FC|>0.15 and adjusted p-value<0.05) either upregulated or downregulated within each cluster in HET^NEG^ (left, cell autonomous changes) and HET^POS^ (right, non-cell-autonomous changes) in the RTT mosaic hippocampus.

We then examined the cell-type specific transcriptional changes in HET^NEG^ and HET^POS^ groups within each cluster, comparing HET^POS^ and HET^NEG^ neurons to WT neurons (**Figure 4E**). First, we found that HET^NEG^ neurons had ∼3X more unique snDEGs (1787) compared to HET^POS^ snDEGs (647) (**Figure 4E**). We also found the magnitude of differential expression (average |log_2_ FC|) was higher in HET^NEG^ DEGs compared to HET^POS^ DEGs across all clusters (**Figure S4E**). In HET^NEG^ neurons, we found snDEGs in both excitatory and inhibitory hippocampal neurons, whereas HET^POS^ snDEGs were almost exclusive to excitatory neurons of the DG, CA1, CA2, and CA3 (**Figure 4E**).

To determine whether changes in HET^POS^ neurons represent a response to the disease or unique response potentially compensating for the disease, we compared the number of snDEGs that overlapped in excitatory clusters between HET^NEG^ and HET^POS^ neurons (**Figure 5A**). We found that ∼25-40% of HET^POS^ snDEGs found in excitatory neurons overlapped with HET^NEG^ snDEGs and these overlapped snDEGs showed a significant strong correlation in DG, CA1, CA2 and CA3 clusters (**Figure 5B**). This indicates that, at least among the shared genes, the majority of non-cell-autonomous responses are changing in the same direction. Interestingly, the CA3-1 cluster had the highest overlap between groups, with ∼79% of HET^POS^ snDEGs overlapping and significantly correlated with HET^NEG^ snDEGs in CA3-1 (**Figure 5B**). We then examined the GO categories of the CA3-1 shared genes (**Figure S5A).** Upregulated genes were enriched in pathways such as membrane potential, potassium ion transport, as well as NAD and glucocorticoid metabolic processes. Downregulated shared genes were enriched in pathways such as synapse assembly, post-synaptic organization, and regulation of synaptic transmission (**Figure S5A**). CA3 neurons are unique compared to other hippocampal cell types in the hippocampus as they form more intraregional connections within CA3 (*46*). While we observed more DEGs in CA1 and DG HET^POS^ neurons, CA3 exhibited a higher degree (CA3: 79%, DG: 25%, CA1: 23%) of concordant changes between HET^POS^ and HET^NEG^ neurons. This pattern of shared transcriptional changes may reflect the extensive local connectivity within CA3, potentially facilitating non-cell-autonomous influences from CA3 MeCP2-to MeCP2+ neurons. Together, these data show that excitatory neurons in the female RTT model are particularly sensitive to the cellular environment created by XCI mosaicism in presymptomatic stages.

**Figure 5.**
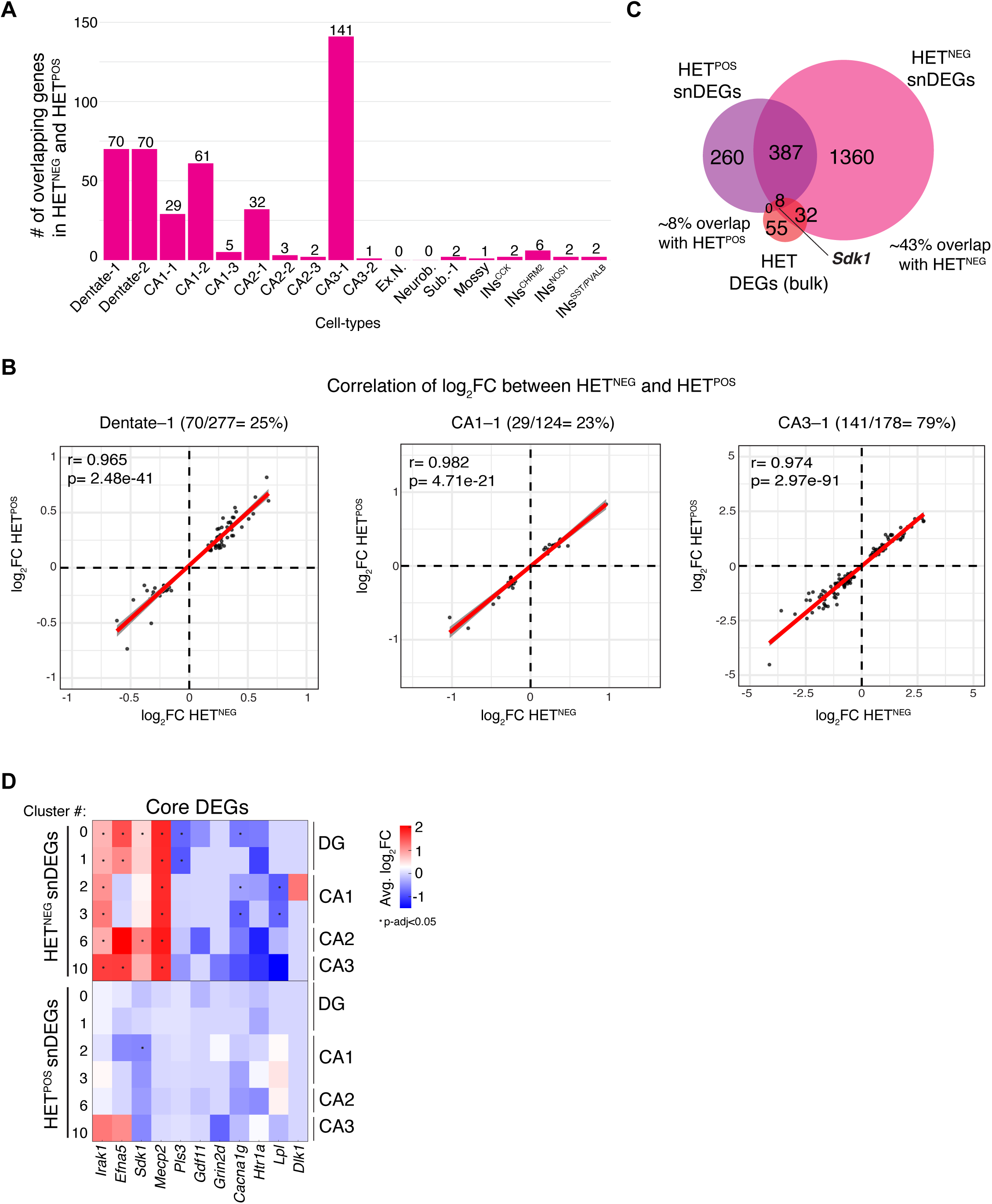
Cell-type specific transcriptional changes in the HET^POS^ and HET^NEG^ cells. **A**) Bar plot showing the number of overlapping snDEGs between neuronal clusters of HET^POS^ and HET^NEG^ samples. **B**) Correlation plots comparing the log_2_FC of snDEGs that overlapped between HET^POS^ and HET^NEG^ samples within the Dentate-1 (DG-1), CA1-1, and CA3-1 clusters. **C**) Venn diagram showing the overlap of bulk RNA-seq DEGs from female HET samples with unique snDEGs from HET^POS^ and HET^NEG^ samples. **D**) Heatmap of log_2_FC of core DEGs found in the HET^POS^ and HET^NEG^ excitatory clusters (* = p-adj<0.05).

### The core molecular disease signature in female RTT is driven by cell-autonomous loss of MeCP2

To examine how the HET^POS^ and HET^NEG^ transcriptional changes in our single-nucleus data contributed to the female bulk RNA-seq disease signature, we overlapped the unique snRNA-seq DEGs from HET^NEG^ and HET^POS^ samples with our HET bulk RNA-seq DEGs (at P45 timepoint) (**Figure 5C**). We found the bulk DEGs overlapped with snDEGs either exclusive to HET^NEG^ (∼43%) or overlapping between HET^NEG^ and HET^POS^ snDEGs (∼8%). Notably, no bulk DEGs overlapped exclusively with HET^POS^ snDEGs (**Figure 5C**). Compared to bulk RNA-seq, ∼7-19X more DEGs were captured in our single-nuclei analysis, further supporting that bulk RNA-seq of mosaic tissue masks many cell-autonomous and non-cell-autonomous molecular changes. We found 11 out of 12 core DEGs (all except *Tpbg*) were captured in our MeCP2-sorted snRNA-seq dataset, likely due to technical differences in 5’ vs 3’ capture of transcript (only 9 out of 12 core DEGs are captured in the previous snRNA-seq dataset). Again, we found that core RTT DEGs were expressed broadly across various cell types in this dataset (**Figure S4D**). We next examined whether the 11 core RTT DEGs were differentially expressed in HET^NEG^ and HET^POS^ excitatory neuron clusters of the DG, CA1, CA2, and CA3 (**Figure 5D**). We found that most core RTT DEGs were specific to HET^NEG^ neuronal clusters and changed in the same direction as our previous bulk and snRNA-seq datasets. Interestingly, *Sdk1* was significantly downregulated in one HET^POS^ cluster of the CA1 region (CA1-1), in the opposite direction from our other datasets, suggesting that molecular compensation of a core DEG occurs in a subpopulation of excitatory neurons (**Figure 5D**). Overexpression of *Sdk1*, which encodes Sidekick-1, a member of the immunoglobulin superfamily of adhesion molecules, has been associated with increased stress susceptibility in the ventral hippocampus (*47*). *Sdk1* also plays a critical role in synapse formation and neuronal connectivity by mediating specific synaptic connections through homophilic adhesion (*48*, *49*). This may explain why *Sdk1* dysregulation in HET^NEG^ neurons could disrupt proper circuit connectivity and affect HET^POS^ neurons. Together, these data show the core molecular signature is largely driven by direct loss of *Mecp2*.

### Cell autonomous gene expression changes in MeCP2- trilaminar neurons

In both of our single-nuclei datasets (male vs female and HET^NEG^ vs HET^POS^), we found 4 subtypes of interneurons (INs^Sst/Pvalb^, INs^Vip/Cck^, INs^Chrm2^ and INs^Nos1^). Of these, INs^Sst/Pvalb^ and INs^Chrm2^ showed the highest number of DEGs. Intriguingly, we observed that one specific interneuron population (INs^Chrm2^) exhibited substantially more snDEGs compared to other interneuron subtypes in both male and female. Based on markers that are expressed from these populations in our dataset and previous literature, we identified this highly affected population as trilaminar interneurons, characterized by high expression of Cholinergic receptor muscarinic 2 (*Chrm2)* while lacking expression of somatostatin *(Sst)*, parvalbumin (*Pvalb),* and other interneuron subtype markers (Cluster 17 in **Figure 6A** and Cluster 21 in **Figure 6B**) (*50–53*). While previous studies have established that MeCP2 function is critical in somatostatin and parvalbumin expressing interneurons, with *Mecp2* knockout in these subtypes alone causing RTT-like phenotypes (*54*), our finding now implicates trilaminar interneurons as another potentially important neuronal population in RTT pathogenesis. Trilaminar neurons are long-range projection interneurons that have been previously described as located in the stratum oriens (SO) and stratum pyramidale (SP) and projecting to the stratum lacunosum-moleculare (SLM) layer of the hippocampus (*52*).

**Figure 6.**
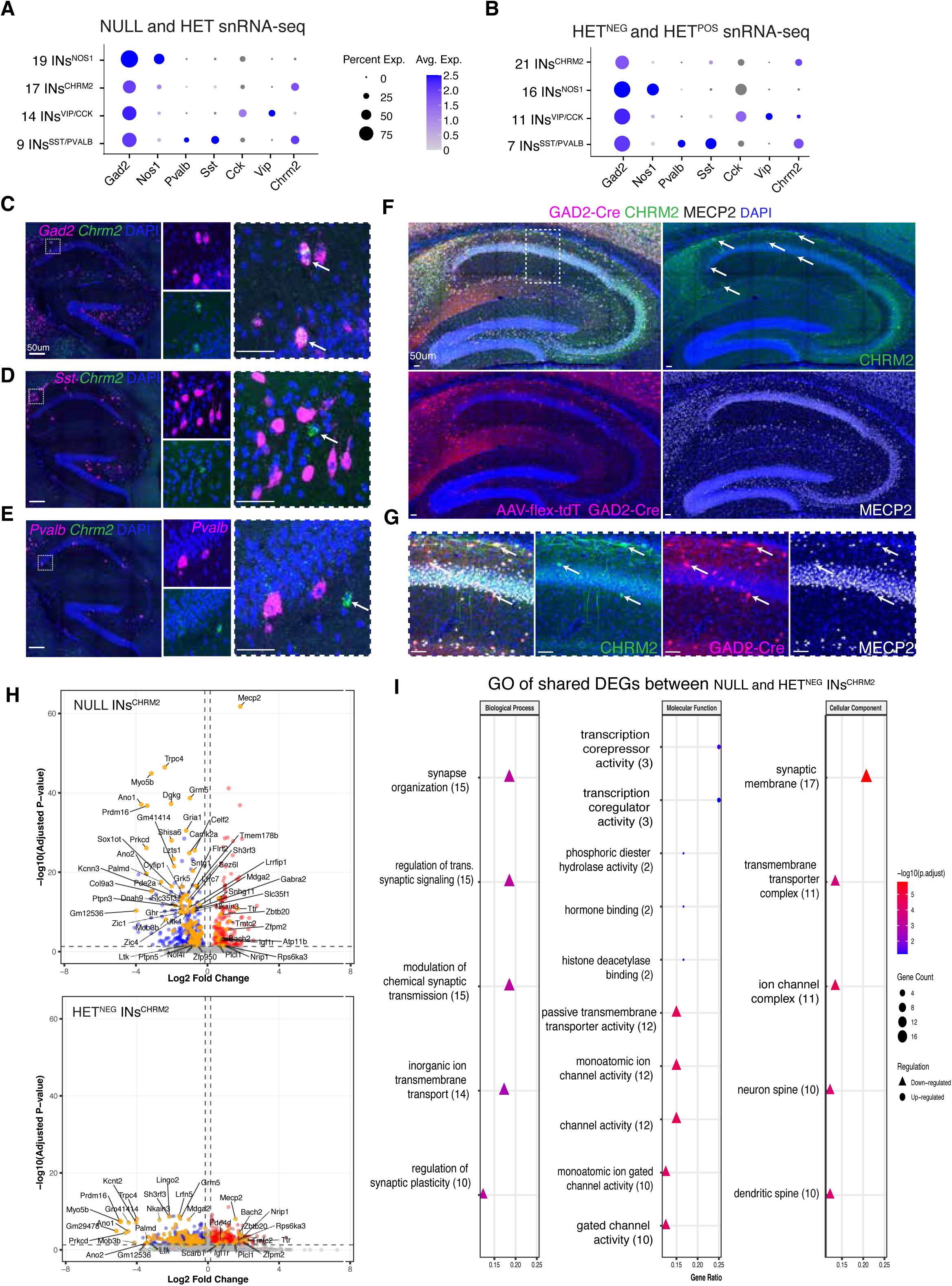
Validation of trilaminar interneuron molecular markers and spatial location in the hippocampus. Dot plots of top interneuron cluster molecular markers in (**A**) both male and female hippocampus datasets and (**B**) female MeCP2-sorted dataset. Dual fluorescent *in situ* images of the hippocampus of wild-type mice showing *Chrm2* transcript (green) co-localizing with (**C**) pan- interneuron marker *Gad2* (magenta), but not with subtype-specific interneuron markers (**D**) *Sst* and **(E)** *Pvalb* (magenta). DAPI stain showing the location of interneurons within the hippocampus. And white arrows indicating *Chrm2* positive neuron within the white bounding box in panel C to E. **F**) Immunofluorescent staining of Chrm2 (green) and MeCP2 (white) in *Gad2-Cre* mice injected with AAV-flex-tdTomato (red) to label interneurons. GAD2 co-localizes with CHRM2 and MECP2 neurons indicated by (**G**) white arrow in bounding box from panel F. **H**) Volcano plot of snDEGs (up-regulated- red, down-regulated-blue) in the trilaminar interneurons (INs^CHRM2^) of the NULL (top) HET^NEG^ (bottom), highlighting shared DEGs in orange between the two datasets. **I**) GO pathways enriched in DEGs that are shared (both upregulated and down downregulated) between trilaminar interneurons populations from NULL and HET^NEG^ samples.

To validate the identity and spatial location of trilaminar interneurons in the hippocampus, we used dual fluorescent *in situ* labeling of *Chrm2* with pan-interneuron marker *Gad2*, as well as two other major interneuron subtypes that also express *Chrm2* (*Sst*, *Pvalb*) (**Figure 6 C-E)**. We confirmed that *Chrm2* colocalizes with *Gad2* and labels a sparse number of interneurons located mainly within the SO and SP layers of CA1-CA3 (**Figure 6C-E**). We identified a subpopulation of *Chrm2* neurons that did not colocalize with *Sst* or *Pvalb* neurons and localized mostly in the CA1 subregion (**Figure 6D- E**). To better visualize the morphology of trilaminar interneurons, we labeled all hippocampal interneurons by injecting *Gad2-Cre* mice with an AAV-flex-tdTomato virus and stained tissue with antibodies against CHRM2 and MeCP2 (**Figure 6F**). We found two morphologically distinct GAD2+/CHRM2+ populations that both colocalized with MeCP2: one located in the upper SO layers with horizontal projecting dendrites, and another in or near the SP layers displaying perpendicular projecting dendrites that spanned the SO and SR hippocampal layers (**Figure 6G**).

Our snRNA-seq data further revealed that trilaminar interneurons with direct loss of MeCP2 (male NULL and HET^NEG^ samples) experience substantial overlap in transcriptional dysregulation (**Figure 6H**). The shared changes between NULL and HET^NEG^ trilaminar interneurons predominantly affect genes involved in neurotransmitter release and synaptic function, with most being downregulated (**Figure 6I**). These findings suggest that trilaminar interneurons are particularly sensitive to direct loss of *Mecp2* compared to other hippocampal interneuron populations early in disease progression in both male and female RTT animal models.

## Discussion

Like other X-linked disorders, RTT has sex-specific differences in clinical presentation due to XCI in females. This creates a mixture of cells expressing either a normal or mutant *MECP2* allele. How this mosaicism contributes molecularly to early RTT pathogenesis remains an open question. In our study, we investigate the mosaic (female) and non-mosaic (male) transcriptional changes in the hippocampus of the RTT mice, using both bulk and single-nucleus RNA sequencing approaches at early timepoints. Combining these techniques, we uncovered shared and distinct cell-type specific disease signatures across both male and female RTT models. Importantly, we identified a set of early core RTT molecular changes common to both males and females, as well as early alteration of the transcriptome in a previously understudied inhibitory neuronal subtype.

### Early core molecular disease signature in the hippocampi of male and female RTT mice

Male and female RTT mice have strikingly different disease progression trajectories (*16*, *19*). Male *Mecp2^-/y^* mice have early onset of phenotypes, including learning and memory deficits, abnormal gait, irregular breathing, hindlimb clasping beginning between 4-6 weeks, and die in young adulthood (∼8 weeks). Female *Mecp2^+/-^* mice have normal survival and display progressive RTT-like neurological phenotypes as they age, typically appearing later in adulthood (∼12-24 weeks) (*16*, *19*). Most studies therefore characterize male mice at a much earlier developmental timepoint than female mice. To exclude molecular changes due to age differences, we compared males and females at the same ages, using early timepoints in disease progression, 4 weeks and 6.5 weeks. While most bulk transcriptional changes were unique between RTT models, with male RTT mice having ∼10X more DEGs than female RTT mice, we found a core molecular disease signature consisting of 12 DEGs altered in the same direction at both timepoints and in both sexes. These core DEGs overlapped with DEGs from a recent study examining the transcriptional changes immediately occurring after the acute loss of MeCP2 in adult *Mecp2* conditional knockout mice (*25*). This suggests that these genes are highly responsive to MeCP2 loss, and they presage disease phenotypes, underscoring their potential contribution to pathogenesis. Eleven of these DEGs were changed in the same direction across all datasets (*Mecp2*, *Pls3*, *Lpl*, *Gdf11*, *Cacna1g*, *Grin2d*, and *Htr1a* downregulated and *Dlk1*, *Sdk1*, *Tpbg*, and *Efna5* upregulated). The only exception is *Irak1* which did not show a significant change upon loss of *Mecp2* in adult mice. Therefore, we propose that these changes appear to be direct consequences of MeCP2 loss (*23*). Consistent with our data, the majority of the core DEGs (*Dlk1*, *Sdk1*, *Efna5*, *Pls3*, *Grin2d*, *Lpl* and *Cacna1g*) are also dysregulated in the same direction in the bulk nuclei RNA-seq data from a study using 6-week-old male mice harboring common RTT missense mutations (T158M and R106W) (*24*, *55*). Moreover, when examining previously published human cortical data (*23*), we found that 5 out of the 12 core DEGs (*EFNA5*, *GDF11*, *HTR1A*, *CACNA1G* and *MECP2*) were also significantly changed in the same direction. Interestingly, several of these core RTT genes are associated with human neurodevelopmental disorders that overlap with RTT phenotypes. For example, mutations in both the N-methyl D-aspartate receptor (NMDAR) subunit *GRIN2D* and calcium voltage-gated channel subunit alpha 1 G (*CACNA1G*) have been shown to cause developmental epileptic encephalopathies (*56*, *57*). The X-linked gene plastin 3 (*PLS3*) has been implicated in severe childhood-onset osteoporosis and scoliosis (*58*, *59*). Together, these findings not only show a core molecular RTT signature across several mouse models but highlight transcriptional changes that precede functional and behavioral changes.

Our results differ from a recent snRNA-seq study that examined the molecular changes over disease progression in cortical tissue from both male and female RTT mice harboring a point mutation at the 5’ transcriptional start site of *Mecp2* (*Mecp2-e1*), which ablates only one *Mecp2* isoform (*22*). These authors did not find the core RTT signature we found shared between male or female RTT models.

Furthermore, they found that peak transcriptional changes occurred in presymptomatic female RTT mice, whereas changes in male RTT mice peaked in symptomatic stages of the disease. Because the adult KO study showed that transcriptomic changes preceded the symptom onsets and correlated with data from constitutive KO male mouse model (R ∼ 0.75) (*25*), we believe the different results are in part driven by the use of the *Mecp2-e1* model where the *Mecp2-e2* isoform is still present and might compensate (*22*).

### Cell-type specific molecular changes in mosaic female RTT hippocampus

In contrast to bulk RNA-seq, we detected robust transcriptional changes in both female and male RTT samples at the single-nucleus level. The shared transcriptional changes in both male and female nuclei were positively correlated with bulk RNA-seq DEGs of *Mecp2^-/y^* mice, suggesting that molecular disease signatures in female RTT tissue (except for the core genes) are largely masked in bulk sequencing due to mosaicism, cellular heterogeneity, and difference in magnitude of gene expression changes. The snRNA-seq data allowed us to uncover cell-type specific changes that are either shared between or unique to male and female RTT models. Most of the shared transcriptional changes between males and females correlated with each other across cell types, except for oligodendrocyte, where snDEGs were upregulated in females and downregulated in males. Previous research has shown that deletion of *Mecp2* specifically in oligodendrocytes did not cause overt RTT phenotypes (*40*) and restoring *Mecp2* in oligodendrocyte lineage results in a minimal rescue.

Altogether, these data indicate that the changes in oligodendrocyte transcription are likely secondary and influenced by the dysfunction of neighboring cells.

### Non-cell-autonomous changes in hippocampal excitatory neurons reveal their responsiveness to the mosaic RTT environment

By sorting MeCP2- and MeCP2+ nuclei for snRNA-seq, we were able to capture cell-type specific direct (cell-autonomous) and indirect (non-cell-autonomous) transcriptional changes in the mosaic *Mecp2^+/-^* hippocampus. Surprisingly, non-cell-autonomous transcriptional changes were only detected in the MeCP2+ excitatory neuronal clusters of the hippocampus. These results add a unique layer of complexity to the effects of E/I imbalance in the female RTT model compared to the male and could explain some of the sex-specific differences observed in *Mecp2* rescue experiments (*38*, *39*). Prior studies have shown that deletion of *Mecp2* in the central nervous system (CNS) can recapitulate RTT phenotypes seen in germline deletion models (*17*). Consistent with that, rescue experiments restoring expression of *Mecp2* in the CNS can reverse most neurological phenotypes (*60*). However, notable differences have been observed between male vs female RTT models in cell-type specific rescue experiments. In female RTT mice, expressing *Mecp2* only in glutamatergic (Vglut2^+^) neurons completely rescued behavioral phenotypes, whereas expressing *Mecp2* only in GABAergic (Viaat^+^) neurons led to only partial improvements (*38*, *39*). In contrast, male mice showed similar improvement in survival and neurological phenotypes when *Mecp2* was in either glutamatergic or GABAergic neurons (*38*, *39*). Considering our molecular findings that MeCP2+ excitatory neurons in the mosaic RTT hippocampus are sensitive to the changes from neighboring MeCP2-neurons, the more effective rescue observed in glutamatergic neurons could be explained by rescuing a larger proportion of dysfunctional neurons early in female RTT mice (**Figure S5C**). Glutamatergic neurons, particularly in the DG, might also be more responsive to the environmental signals or changes in network activity, since this region is a niche for neurogenesis in the adult brain and is primed to respond to external cues (*61*).

Our findings align with previous studies that have found non-cell-autonomous transcriptional changes in MeCP2+ cells of RTT female cortical neurons (*22–24*). Work from *Sharifi et al*. showed large non-cell-autonomous changes in both excitatory and inhibitory neurons in the cortex at the earliest time point (P30), which by P60 were largely restricted to glutamatergic neurons (*22*). *Renthal and Boxer et al*. focused their analysis of non-cell-autonomous changes exclusively in cortical glutamatergic neurons and found non-cell-autonomous changes in the 12-20-week-old females RTT mice (*23*).

*Johnson, Zhao and Fasolino et al.* found non-cell-autonomous changes in MeCP2+ cells in female mice modeling common RTT-causing mutations (T158M and R106W); however, they did not examine cell-type specificity further (*24*). Our study focuses on both excitatory and inhibitory neurons and reveals regional cell-type specific differences, with a non-cell-autonomous influence particularly on excitatory neurons of the hippocampus early in disease pathogenesis. Consistent with our transcriptional findings, previous studies examining morphological changes of MeCP2+ cells in older (5-21 month) female *Mecp2^+/-^* mice compared to control neurons found significant decreases in soma size, nuclear size, and spine density (*44*, *62*). Although functional characterization of basal electrophysiological properties of MeCP2+ neurons in *Mecp2^+/-^* mice have not found clear differences compared to controls(*26*, *27*), our findings suggest that mosaicism within the female hippocampus influence the transcriptome of glutamatergic excitatory neurons early in the disease process and could influence functional deficits via both direct and indirect molecular mechanisms.

### Transcriptional changes are enriched in trilaminar interneurons across RTT models

Our snRNA-seq studies uncovered a particular subcluster of interneurons that displayed ∼3-4x more significant transcriptional changes upon loss of *Mecp2* compared to other interneuron populations.

The molecular changes in this interneuron subcluster were striking given they were found across three distinct comparisons in both male and female RTT and are unique to MeCP2- nuclei of female RTT samples. These interneurons have molecular characteristics of hippocampal trilaminar interneurons, including high expression of *Chrm2* while lacking expression of *Sst* and *Pvalb*(*50*), consistent with a previous study that profiled different subclasses of CA1 inhibitory neurons (*63*).

Although trilaminar hippocampal interneurons have not been extensively characterized, studies have shown they localize to the stratum oriens and pyramidale layers and they input onto somatostatin interneurons in the OLM (SST+ OLM) (*50–53*, *63*, *64*). This connection is particularly intriguing because previous research has shown that MeCP2-deficient SOM+ OLM interneurons in *Mecp2^+/-^* females are weakly recruited into associative memory ensemble, resulting in larger CA1-excitatory neuronal engrams and deficits in long-term memory recall (*27*). Moreover, *Chrm2* KO mice have significant deficits in memory and synaptic plasticity (*65*), suggesting that disruption of cholinergic signaling in these neurons may contribute to cognitive dysfunction. The disproportionate vulnerability of these neurons to MeCP2 loss, given the extensive transcriptional dysregulation of genes involved in neurotransmission and synaptic function, highlights a potentially important mechanism underlying early RTT pathogenesis. It will be crucial to investigate how this specific neuronal population contributes to circuit dysfunction in RTT and whether therapeutically targeting these neurons could ameliorate cognitive symptoms of the disorder.

In summary, our study revealed: 1) a core set of genes consistently dysregulated across different timepoints in both male and female RTT mouse models, indicating shared and early molecular mechanisms in RTT pathogenesis; 2) single-nucleus RNA sequencing uncovered significantly more transcriptional alterations compared to bulk RNA sequencing in female RTT mice, highlighting the importance of cellular resolution in studying mosaic disorders; 3) notable cell-type specific differences between male and female models in spite of many shared gene expression changes; 4) distinct non-cell-autonomous effects primarily affecting excitatory neurons in the mosaic female RTT hippocampus and 5) extensive transcriptional changes unique to the MeCP2 negative trilaminar neurons in both male and female RTT mice. Together, these data reveal how female mosaicism influences distinct molecular pathways in the hippocampus, providing crucial insights that might extend beyond Rett syndrome to other X-linked neurodevelopmental disorders such as DDX3X syndrome and CDKL5 deficiency disorder. These findings underscore the fundamental importance of considering sex-specific molecular mechanisms in disease pathogenesis and highlight the critical need for therapeutic strategies that account for the unique biology of female mosaicism across X-linked neurological conditions.

## Materials and Methods

### Animals

Mice were maintained on a C57BL/6 background on a 14-h:10-h light:dark cycle at 68 to 72°F and 30 to 70% humidity and fed standard mouse chow and water *ad libitum.* Up to five mice were housed per cage. *Mecp2^-/y^* and *Mecp2^+/-^* mice were maintained by crossing *Mecp2^+/-^* females to wild-type C57BL/6J males (*16*). Experimental mice were obtained through crossing *Mecp2^+/-^* to C57BL/6J from the Jackson Laboratory. Homozygous male and female Gad2-IRES-Cre knock-in C57BL/6J mice (*Gad2^tm2(cre)Zjh^*) were obtained from Jax (Strain #028867) and homozygous mice were bred together and genotyped according to the Jax protocol. The Baylor College of Medicine Institutional Animal Care and Use Committee approved all research and animal care procedures.

### Bulk RNA sequencing

Hippocampi were dissected from P28 or P45 mice and flash-frozen in liquid nitrogen (n = 4 per genotype). Tissues were homogenized using a TissueLyser II (Qiagen, #85300) with a customized program (30.0 Hz, 30 s × 2). Total RNA was extracted immediately using the RNeasy Mini Kit (Qiagen, #75106) according to the manufacturer’s instructions, including on-column DNase I treatment (Qiagen, #79254) to remove genomic DNA. RNA quality was assessed by Genewiz using TapeStation and Qubit, followed by library preparation and sequencing with rRNA depletion method for mRNA with UMI controls. Libraries were sequenced at ∼50 million 150-bp paired-end reads per sample for transcriptomic profiling. STAR v2.7.9a was used to align trimmed reads to the Mus musculus genome (mouse_mm10v23 and gencode.vM23.primary_assembly.annotation.gtf for annotation) to obtain read counts with default parameters (*66*). Default STAR parameters were used other than –sjdbOverhang 149. The alignment rates for all samples were above 80%. The read count matrix was used as input for differential gene expression analysis with DESeq2 (v1.40.2) (*67*). Genes with a mean count <10 were filtered out to remove lowly expressed genes. Differentially expressed genes (DEGs) were identified using an adjusted p-value threshold of <0.05 and abs(log2FC) > 0.15. Overlap of the 12 core RTT DEGs with acute knockout data was done using DEGs from the 6 week post MeCP2 deletion (*25*). A hypergeometric test was performed using 1072 DEGs in adult KO and 20,000 for background genes, observed overlap is was 11 genes.

### Tissue collection and nuclei isolation

Animals were decapitated after anesthetization with 3% isoflurane. The hippocampi of mice were dissected on ice, flash frozen in liquid nitrogen, and stored at -80 °C until further use. The nuclei isolation protocol was adapted from 10x Genomics for adult brain tissue. Briefly, on the day of nuclei isolation, frozen hippocampi were directly homogenized in ice cold lysis buffer (10mM Tris-HCl pH7.4, 10mM NaCl, 3mM MgCl2, 0.1% IGEPAL CA-630 from Sigma-Aldrich) and incubated on ice for 10 min. The tissue lysate was passed through a 30-μm MACS SmartStrainer filter to remove larger debris and then centrifuged at 500 x g for 5 min at 4°C. Supernatant was removed and the nuclei pellet was washed with wash buffer (1x Dulbecco’s phosphate-buffered saline (DPBS) with 1% BSA (ThermoScientific #37525) and then centrifuged at 500xg for 5 min at 4°C. A total of 2 washes were performed. For samples requiring antibody staining, MeCP2-conjugated PE antibody (Cell Signaling Technology #34113S) was used and incubated at 1:1000 dilution at 4°C for 30 min followed by 2x washes. The final nuclei suspension was checked under Thermo Fisher Countless II using Trypan blue to ensure the particle concentration was below 1 x 10^7 to avoid clogging on the SONY SH800s machine.

### Nuclei sorting

Nuclei suspensions were stained with DRAQ5 fluorescent probe (1:1000 dilution) for 5 minutes before sorting. The forward scatter (FSC) threshold was set at 1.50% with an FSC gain of 11, and the back scatter (BSC) gain was adjusted to 30.5%. Nuclei gates were first determined by passing samples through a gate (BSC-A vs FSC-A) to filter out debris using control nuclei without DRAQ5 staining, then through two single nucleus gates (FSC-H vs FSC-A and BSC-H vs BSC-W). For experiments separating HET^POS^ and HET^NEG^ group, the HET^NEG^ gate was determined using stained nuclei from NULL male. Prior to sorting, the 1.5 mL collection tube was coated with 1% bovine serum albumin (BSA) and filled with 50 μL of single cell sorting buffer (1X DPBS with 1% BSA). 100,000 - 150,000 nuclei were collected for each sample. For HET-sorted samples, HET^POS^ and HET^NEG^ group were sorted into separate tubes. After sorting, nuclei were spun down at 1500 × g for 10 min at 4°C to recover all nuclei and then resuspended to roughly a concentration of 1000 nuclei/μL. 10,000 to 12,000 nuclei were loaded on the 10X Chip.

### Library construction and sequencing

The cDNA libraries were constructed by 10x Genomics 3’ v3.1 or 5’ HT Reagent Kits v2 (Dual index) following the user guide. Briefly, a total volume of 77.4uL of nuclei suspension (10000 nuclei targeted) were mixed with reverse transcription master mix before loading into Chromium Chip G/N. Droplets containing nuclei, reverse transcription reagents, and barcoded gel beads were generated by Chromium X Controller. The first strand cDNA was then amplified and checked using TapeStation with HS D5000 ScreenTape to assess cDNA amplification. Then the cDNA libraries were fragmented, size selected using SPRI beads and ligated with sequencing adapters and sample indices. The cDNA libraries were sequenced by Illumina NovaSeq 6000 using S4 flow cell chemistry.

### Data pre-processing

The reads were aligned to 10X Genomics mouse reference genome mm10 (2020-A). The alignment and quantification of unique molecular identifiers (UMI) were performed on 10X Genomics Cloud by the Cell Ranger pipeline v7.0.1 with default parameters including intronic reads mapping.

We performed quality control for the single-nucleus RNA-seq data using the R package Seurat (v5.0.0). Specifically, we used the following criteria to identify low-quality nuclei: 1. UMI counts < 500 or UMI counts > 30,000; 2. number of genes < 500 or number of genes > 6000; 3. mitochondrial gene expression ratio > 2%. We filtered out all low-quality nuclei. For doublet removal, we used the R package scDblFinder (v1.12.0) to detect potential doublets and removed all nuclei classified as doublets. Nuclei from different samples were integrated with batch effect correction using the R package harmony (v1.2.0) (*68*). The integrated data were analyzed following the Seurat workflow. Specifically, we (1) normalized the raw counts (normalization.method = "LogNormalize"), (2) selected the top 2,000 highly variable genes, and scaled the data based on these highly variable genes, (3) used the selected highly variable genes to calculate the first 50 principal components, (4) clustered nuclei using the FindNeighbors function (dims = 1:20) and the FindClusters function (resolution = 0.5), and (5) reduced the dimension of the data through UMAP (dims = 1:20).

### Cell annotation

The top marker genes of different clusters were identified using the FindAllMarkers function (only.pos = TRUE, min.pct = 0.25, logfc.threshold = 0.25) from the R package Seurat (v5.0.0). Cell type identity was determined by comparing these top marker genes to published reference datasets (*42*, *43*, *50*, *63*, *69*, *70*). Cell type composition testing was performed using propeller function from R package speckle (*71*), taking into account individual samples within each experimental group.

### Identification of differentially expressed genes (DEGs) in snRNA-seq

The DEG analysis was performed using the Seurat function FindMarkers (min.pct = 0.1) with the MAST method (v1.24.1). With the output from FindMarkers, DEGs were defined using the following criteria: 1. The absolute value of average log2 fold change > 0.15. 2. The adjusted p-value < 0.05. Upset plots for DEG numbers among different comparisons were generated using the R package ComplexUpset (v1.3.5). Unique snDEGs were defined as genes that were differentially expressed in one or more clusters, regardless of the number of clusters in which they appeared.

### Pathway Analysis

Gene Ontology (GO) pathway enrichment analysis was performed using the R package clusterProfiler (v4.6.2) with the function enrichGO (OrgDb = org.Mm.eg.db (v3.16.0), pAdjustMethod = "BH", pvalueCutoff = 0.05, qvalueCutoff = 0.05). The up-regulated (log2FoldChange > 0) and down-regulated (log2FoldChange < 0) DEGs were used as input to enrich up-regulated and down-regulated pathways.

### Trilaminar interneuron labelling

P0 pups from homozygous *Gad2^tm2(cre)Zjh^* breeding received bilateral intracerebroventricular (ICV) injections with AAV9-FLEX-tdTomato virus obtained from Addgene (Catalog #28306-AAV9), using a 1ul volume with 1.5e^10 viral particles per hemisphere. Mice were harvested 25 days later for downstream processing. Following viral injection, mice at age P25 were completely anesthetized by giving an intraperitoneal injection of Rodent Combo III, 1.5ml/kg, then perfused with cold 30ml of 1X PBS and 30ml of 4% PFA. Brains were extracted and incubated in 4% PFA overnight. Brains were washed with cold 1X PBS and then incubated in 30% sucrose with 0.01% sodium azide solution for at least 48hrs until sectioning. A Leica SM2010R sliding microtome at -25°C stage temperature was used to section right hemisphere along the sagittal plane at 60um thickness. Free floating serial sections were stored in 1X PBS + 0.01% sodium azide at 4°C until further processing. Sections were washed three times with PBS and incubated in permeabilization buffer (0.3% Triton-X-100 in PBS) for 5 min at room temperature. Sections were placed in blocking buffer (0.3% Triton-X, 3% normal donkey serum) for 30 min at room temperature and then incubated with primary antibodies (rtChrm2 MAB367 1:500, rbMECP2 CST34113S 1:500) with 1% BSA in blocking buffer overnight, rocking at 4°C. Sections were washed three times with PBS then incubated with secondary antibodies at 1:1000 dilutions (donkey anti-rabbit 647 from Jackson ImmunoResearch 711-605-152 and goat anti-rat 488 from Thermo Scientific A-11006) in blocking buffer with 1% BSA for 1 hr at room temperature. After final washes with PBS, sections were incubated for 10 min in 5ug/ml DAPI solution then mounted onto slides and allowed to dry. Coverslips were mounted using Prolong Gold mounting media and slides allowed to dry overnight. Stained sections were imaged on a Leica SP8X confocal microscope using 20X oil objective.

### Dual-color fluorescence RNA in situ hybridization

Fluorescent in situ hybridization was performed by the RNA In Situ Hybridization Core at BCM using previously described methods (*72*). Briefly, fresh frozen adult mouse tissue was used and sectioned along the sagittal plane at 25um. Digoxigenin (DIG)-labeled, or fluorescein (FITC)-labeled mRNA antisense probes were generated for *Gad2*, *Chrm2*, *Sst*, and *Pvalb* from Allen Brain Atlas sequences (https://mouse.brain-map.org). The probe templates were generated by PCR using reverse-transcribed mouse cDNA as template and DIG or FITC RNA labeling kits from Roche (Sigma). Slides were imaged on a Leica SP8X confocal microscope using 20X objective.

## Supplemental Figure Legends

**Supplemental Figure 1.**
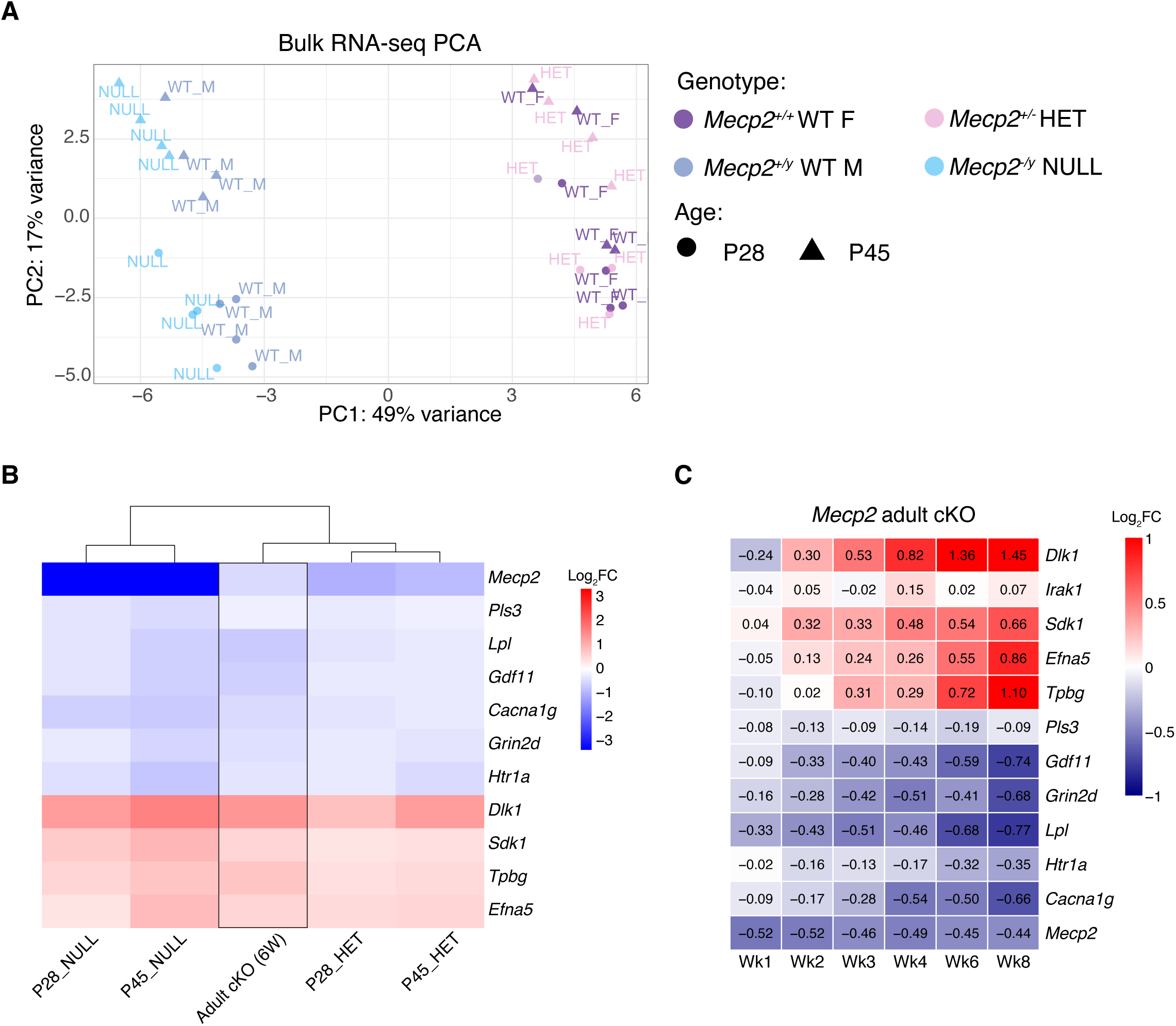
PCA of bulk RNA-seq samples and overlap with *Mecp2* adult conditional (adult cKO) knockout gene expression data. **A**) Principal component analysis (PCA) of all 32 hippocampal samples from male and female WT, NULL, and HET samples at two timepoints. **B**) Heatmap comparing the log2FC of 11 core DEGs found in the hippocampal adult *Mecp2* cKO bulk RNA-seq to our dataset. **C**) Heatmap of log2FC of all core DEGs across the time course of adult *Mecp2* cKO bulk RNA-seq at week (wk) 1, 2, 3, 4, 6, and 8 post-deletion of *Mecp2*.

**Supplemental Figure 2.**
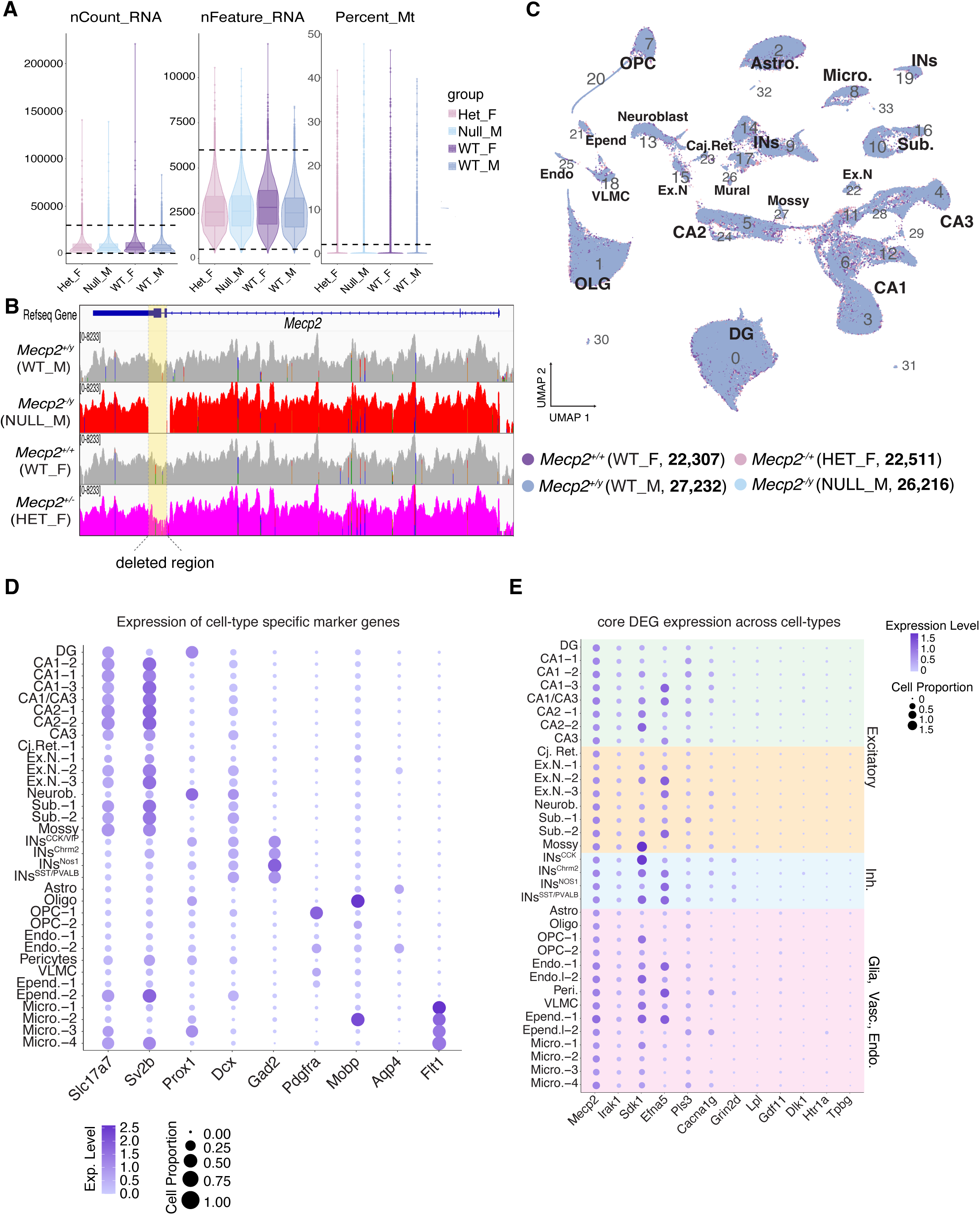
snRNA-seq quality control measures across genotypes and cell type annotation of clusters. **A**) Distribution of key quality control metrics across genotype groups. Violin plots show distribution of unique molecular identifiers (UMIs) (nCount_RNA), genes detected (nFeature_RNA), and percent mitochondria (Percent_Mt) with the black dotted lines identifying the cutoff quality control values (see methods). Following quality control filtering, there were no significant difference between genotype groups (Kruskal-Wallis test, p = 0.74). **B)** Track files of all *Mecp2* reads from the single-nuclei dataset separated by genotype, showing that no reads map to the deleted exons 3 and 4 (yellow highlight) of *Mecp2*^-/y^ mice and reads in the exons are reduced in *Mecp2*^+/-^ mice. **C**) UMAP showing nuclei labelled by genotype. **D**) Bubble plot showing expression of top cell type marker genes within each cluster. **E**) Bubble plot showing expression of core DEGs across cell types.

**Supplemental Figure 3.**
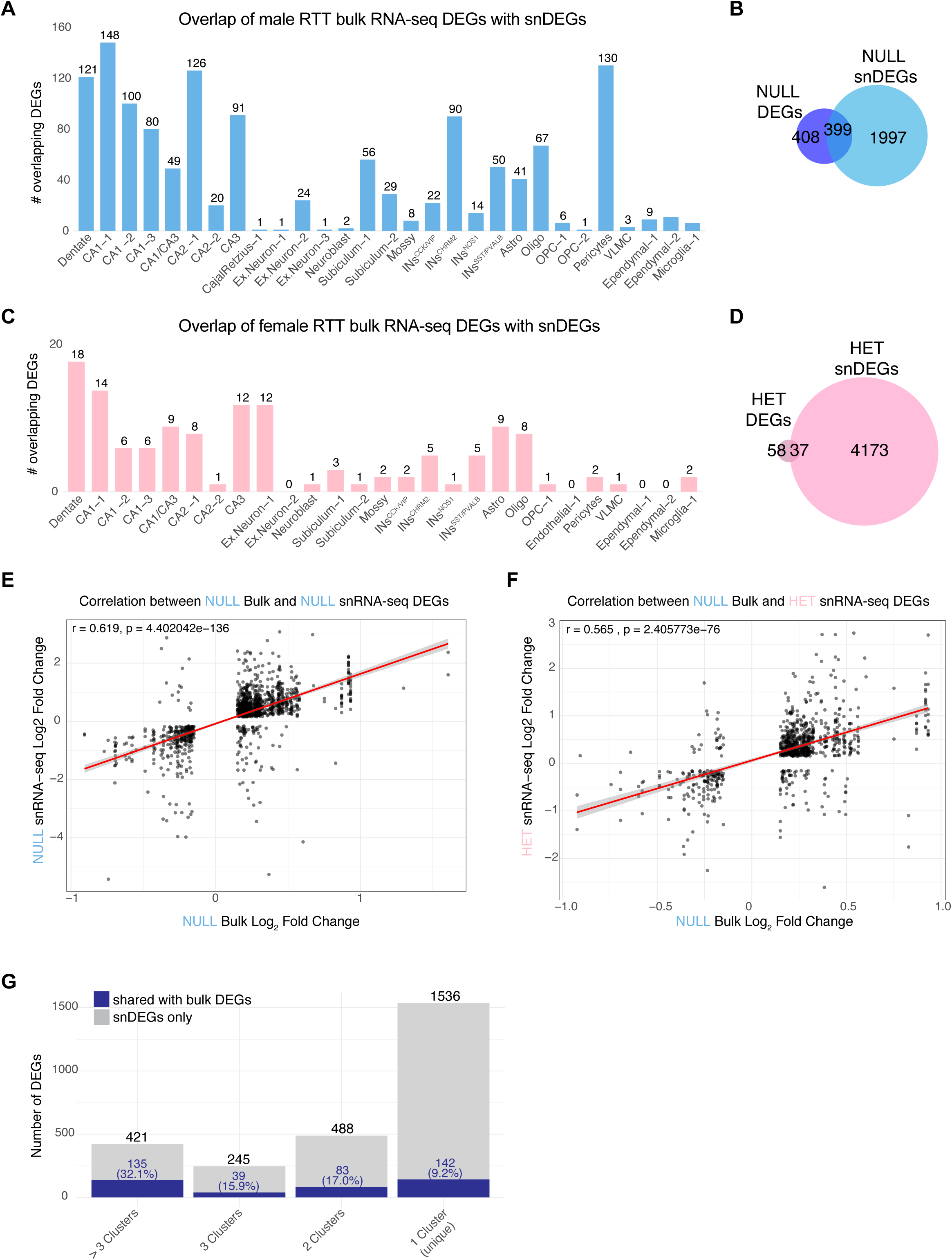
C**e**ll**-type specific overlap of bulk RNA-seq DEGs with snDEGs in hippocampus of male and female RTT mice. A**) Bar charts showing the number of overlapping bulk RNA-seq DEGs with snDEGs across cell types in male (top panel, blue) or (**C**) female (bottom panel, pink) RTT samples. Venn diagram showing the overlap of unique snDEGs with bulk DEGs in (**B**) male and (**D**) female samples. Correlation plots of log_2_FC between male NULL bulk RNA-seq with snRNA-seq DEGs from (**E**) male NULL or (**F**) female HET samples. **G**) Bar plot showing the number of male snDEGs that are unique to one cell type (1 cluster) or shared across multiple clusters (gray bars), with blue bars indicating overlap with male bulk DEGs (P45) and percentages showing overlap rates. Data from male snDEGs, female shows similar pattern (data not shown).

**Supplemental Figure 4.**
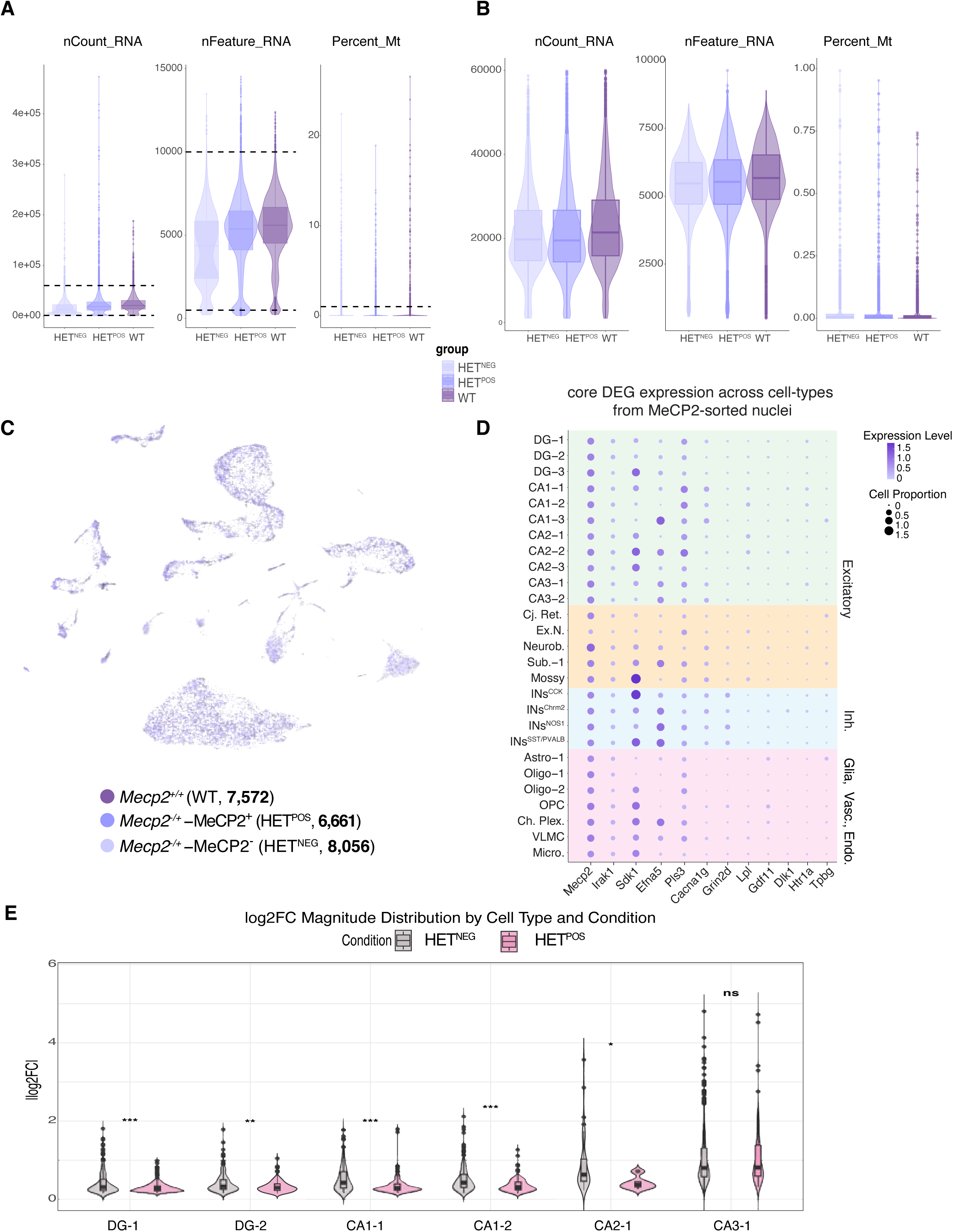
s**n**RNA**-seq quality control measures within the hippocampus of female RTT mice. A)** Violin plots show distribution of unique molecular identifiers (UMIs) (nCount_RNA), genes detected (nFeature_RNA), and percent mitochondria (Percent_Mt) with the black dotted lines identifying the cutoff quality control values and **B)** same QC plot post-filtering for neuronal population. Following filtering, nCount_RNA shows no significant difference between genotype groups (Kruskal-Wallis test, p = 0.18). **C)** UMAP showing nuclei labelled by sample groups. **D**) Bubble plot showing expression of core DEGs across cell types. **E)** Violin plots showing the distribution of absolute log2 fold change (|log2FC|) values for snDEGs in HET^NEG^ (MeCP2-, gray) and HET^POS^ (MeCP2+, pink) neurons compared to WT controls across different hippocampal neuronal subtypes. Each point represents an individual snDEG. HET^NEG^ neurons consistently show higher magnitude of transcriptional changes compared to HET^POS^ neurons across most cell types. Statistical significance was determined by Wilcoxon rank-sum test. *p < 0.05, **p < 0.01, ***p < 0.001, ns = not significant.

**Supplemental Figure 5.**
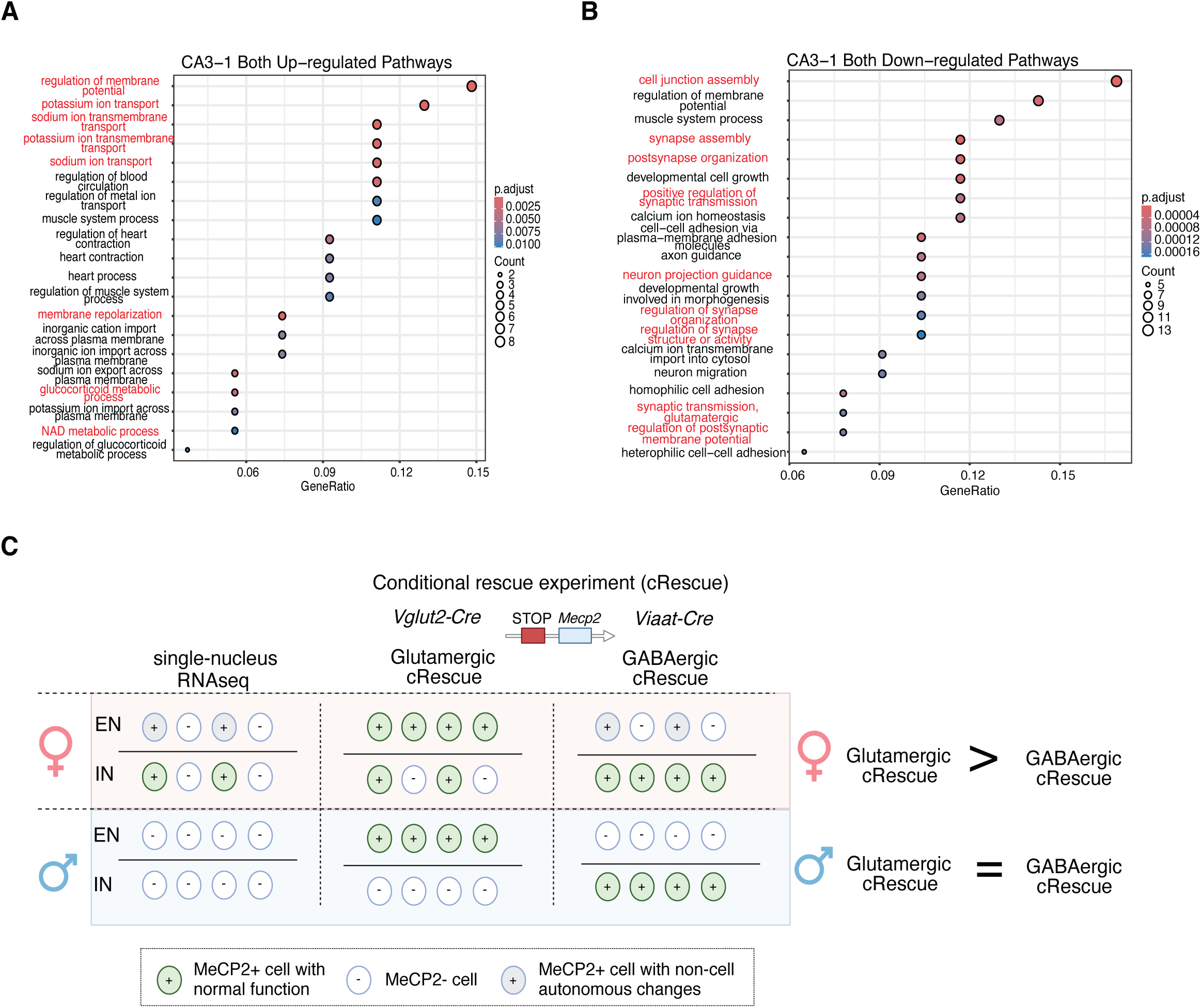
G**O analysis of shared CA3 snDEGs in mosaic RTT and graphic illustration of cell-type specific scenarios as related to conditional rescue experiments in male and female RTT mice**. GO pathways enriched in shared genes that are (**A**) upregulated or (**B**) downregulated in both HET^POS^ and HET^NEG^ in the CA3-1 cluster. **C**) A working model placing our snRNA-seq findings in the context of previous sex-specific differences observed in conditional rescue experiments in male vs female RTT mice. In the mosaic female RTT mice, re-expressing *Mecp2* in glutamatergic, excitatory neurons rescued neurological phenotypes better than re-expressing *Mecp2* in inhibitory, GABAergic neurons. Our snRNA-seq results show that glutamatergic neurons are particularly disrupted in the mosaic RTT brain, where both MeCP2+ and MeCP2- cells display altered transcriptional responses, compared to the GABAergic neurons where only MeCP2- cells are affected. Therefore, restoring *Mecp2* in glutamatergic neurons rescues both non-cell-autonomous and cell-autonomous changes (i.e. a larger proportion of neurons) than only rescuing cell-autonomous changes in GABAergic interneurons. In male RTT models, re-expressing *Mecp2* in either glutamatergic or GABAergic neurons had similar levels of rescue, particularly on survival. Graphical illustration created using BioRender.com.

## Acknowledgements

We would like to thank to Dr. Harini Tirumala, Dr. Amanda Engstrom, Dr. Rebecca Meyer Schuman, Dr. Joshua Brenner, Cole Deisseroth, Siming Zhong and Rui Zhang for their valuable input and critical feedback on the manuscript. We also thank Dr. Hari Yalamanchili and Dr. Rambabu Majji for developing and implementing the alignment pipeline for bulk RNA-seq data.

Special thanks to Dr. Jessica Butts for providing training on the 10X Genomics platform. We would like to thank Yaling Sun, Kailey Xia, and Elizabeth Hai-Yen Chu for assisting with the mouse colony maintenance to make our work possible. Their collective expertise and support were essential to the completion of this work. This work was supported by the National Institute of Neurological Disorders and Stroke (NINDS) (R01NS057819 to HYZ and F32N122920-01A1 to AGA) and the Howard Hughes Medical Institute (to HYZ). The project described was supported in part by the RNA In Situ Hybridization Core facility at Baylor College of Medicine, which is supported by a Shared Instrumentation grant from the NIH (1S10OD016167) and the NIH IDDRC grant P50 HD103555 from the Eunice Kennedy Shriver National Institute of Child Health & Human Development. The content is solely the responsibility of the authors and does not necessarily represent the official views of the Eunice Kennedy Shriver National Institute of Child Health & Human Development or the National Institutes of Health.

## Author contributions

YL and AGA contributed equally as co-first authors of this work and have permission to place their name in the first author position on their respective CVs. YL, AGA, and HYZ conceptualized the project. YL, AGA, JPR and SRW generated or helped generate the snRNA-seq data. GQ and YL processed the snRNA-seq data. YL, AGA, GQ annotated and analyzed the snRNA-seq data. YL generated and processed the bulk RNA-seq data. YL and AGA analyzed the bulk RNA-seq data. YL, AGA, GQ and HYZ interpreted data. AGA performed the interneuron staining and imaging experiments. AGA wrote initial draft of manuscript. YL, AGA, GQ and HYZ edited the manuscript. All authors reviewed and provided comments on the manuscript.

